# Bone Marrow Polyethylene Microplastics Favour Tet2-mutant Haematopoiesis and Aggravate Atherosclerosis

**DOI:** 10.64898/2026.11.20.767985

**Authors:** Xinning Guo, Yuyan Lyu, Yongdan Yang, Songmao Wang, Haibo Zhang, Xin Wang, Gangjian Qin, Qing Rex Lyu

## Abstract

We investigated whether polyethylene microplastics (PE-MPs) in bone marrow influence clonal haematopoiesis and atherosclerosis. Pyrolysis-gas chromatography/mass spectrometry detected polyethylene in bone marrow aspirates from 27 of 31 adults (87.1%). In *Ldlr*^−/−^ mice competitively transplanted with wild-type and *Tet2*^+/−^ marrow, intratibial PE-MPs exposure increased atherosclerotic lesion area in the aortic arch and root, the relative contribution of *Tet2*^+/−^ donor cells, and the abundance of mutant-derived myeloid cells. In a separate cohort reconstituted with *Tet2*^+/−^ marrow alone, single-cell RNA sequencing revealed transcriptional signatures of granulocyte-monocyte lineage priming and cellular stress in haematopoietic stem and multipotent progenitor cells following PE-MPs exposure. These findings suggest that marrow PE-MPs exposure can modify *Tet2*-mutant clonal haematopoiesis and exacerbate atherosclerosis in mice.

## Introduction

Microplastics have been detected in human bone marrow and atherosclerotic plaques, raising concerns about their cardiovascular consequences ^1, 2^. Clonal haematopoiesis (CH), particularly involving TET2 mutations, is associated with increased cardiovascular risk and accelerated atherosclerosis ^3, 4 5^. However, whether bone marrow microplastics influence mutant haematopoiesis and atherosclerosis remains unknown. We investigated the effects of local bone marrow polyethylene microplastics (PE-MPs) exposure on the competitive contribution of *Tet2*-mutant haematopoietic cells and atherosclerotic lesion development.

## Methods

Bone marrow aspirates (∼1mL) were collected in glass containers from adults (≥18 years) underwent clinically indicated aspiration. Participants had capacity to consent independently and gave written informed consent for research use of residual aspirates and deidentified clinical data. Exclusions were pregnancy or lactation, incarceration, myeloid neoplasms, prior hematopoietic stem cell transplantation, and recent intensive chemotherapy, radiotherapy, or other treatments likely to alter the marrow microenvironment or clonal hematopoiesis assessment. After sequential solvent extraction of dried aspirates, 13 polymers were quantified by pyrolysis-gas chromatography/mass spectrometry (Py-GC/MS) using polymer-specific pyrolysis products, characteristic ions, and external calibration. Each batch included instrument, reagent, procedural, and sampling blanks. Polymer-specific blank masses were subtracted before normalization to dry tissue mass. Corrected values below sample-specific detection limits were recorded as undetected. The Medical Research Ethics Review Committee of the First Affiliated Hospital of Chongqing Medical University approved the study (ZZ2026-0827-01).

Male *Ldlr*^−/−^ mice aged 6∼8 weeks received 8 Gy total-body irradiation and unilateral intratibial transplantation of 5×10^6^ marrow cells in 10 μL. The competitive cohort received a 1:1 mixture of CD45.1 WT and CD45.2 *Tet2*^+/−^ marrow, and the independent noncompetitive cohort received *Tet2*^+^/^−^ marrow for transcriptional profiling. After four weeks, mice were randomized to a single bilateral intratibial injection of 20μm polyethylene microplastics (PE-MPs) (1μg in 10 μL PBS per tibia) or phosphate-buffered saline (PBS) and fed a high-fat, high-cholesterol diet (D12108; Research Diets) for 16 weeks. Atherosclerosis was quantified by *en face* and aortic-root serial Oil Red O staining. Bone marrow scRNA-seq included three control and five PE-MPs-exposed mice. Donor chimerism was measured among viable CD45^+^ cells; GMPs and mature myeloid populations were identified using Lineage/Sca-1/c-Kit/CD34/CD16/32 and CD11b/Ly6C/Ly6G/F4/80 panels, respectively. Absolute counts were calculated from viable-cell counts and sequential gated frequencies. Outcome assessment was blinded. The Yi Shang Biotech Institutional Animal Care and Use Committee approved all animal procedures (IACUC-2024-Mi-051).

## Results

We first quantified 13 polymers in residual bone marrow aspirates from 31 adults (19∼85 years) undergoing clinically indicated aspiration. Samples were collected in glass containers and analysed using pyrolysis-gas chromatography/mass spectrometry (Py-GC/MS), with instrument, reagent, procedural, and sampling blanks. Polymer-specific blank values were subtracted before normalization to dry tissue mass, and corrected values below sample-specific detection limits were classified as undetected. Polyethylene (PE) was detected in 27 of 31 samples (87.1%), with a mean concentration of 8.42±7.45 µg/g (**Figure 1A**). Given its frequent detection in human bone marrow and reported presence in atherosclerotic plaques ^1, 2^, PE was selected for experimental investigation.

**Figure 1.**
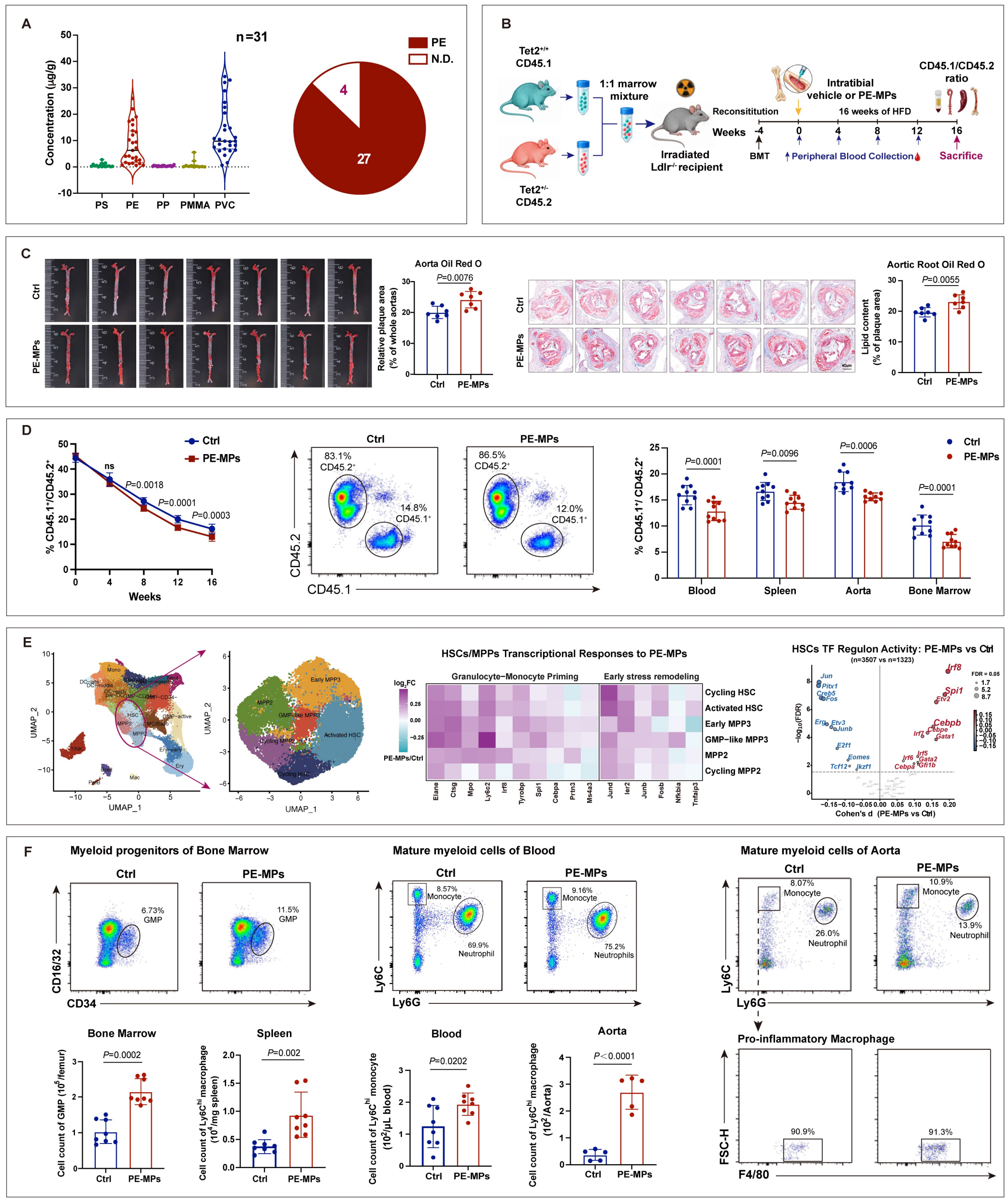
Bone marrow polyethylene microplastics exposure increases atherosclerosis and alters *Tet2*-mutant haematopoietic competition. **A**. Quantification of polymer concentrations in human bone marrow aspirates by pyrolysis-gas chromatography/mass spectrometry (Py-GC/MS; n=31). Each dot represents one participant. Polyethylene was detected in 27 samples (87.1%). ND, not detected. **B**. Experimental design of competitive bone marrow transplantation. Irradiated *Ldlr*^−/−^ recipients received equal numbers of CD45.1 wild-type (WT) and CD45.2 *Tet2*^+/−^ donor marrow cells, followed by intratibial polyethylene microplastics (PE-MPs) or phosphate-buffered saline (PBS) and 16 weeks of high-fat feeding. **C**. Representative *en face* aortic arch and aortic-root Oil Red O staining and quantification of atherosclerotic lesion area (n=7 mice per group). Aortic-root measurements represent the mean of nine serial sections per mouse. **D**. Longitudinal peripheral-blood WT/*Tet2*^+/−^ donor-cell ratios and corresponding ratios in blood, bone marrow, spleen, and aorta at week 16 (blood, spleen, bone marrow, n=10; aorta, n=9 per group). **E**. Single-cell RNA sequencing of bone marrow from a separate *Tet2*^+/−^-only transplantation cohort (PBS, n=3; PE-MPs, n=5). UMAP plots, HSCs/MPPs subclusters, selected lineage-priming and stress-response genes, and inferred myeloid-associated transcription-factor regulon activities. **F**. Representative flow-cytometry plots and quantification of *Tet2*^+/−^-derived bone marrow granulocyte– monocyte progenitors, peripheral-blood Ly6C^hi^ monocytes, and splenic and aortic Ly6C^hi^ macrophage populations (marrow, blood, and spleen, n=8; aorta, n=5 per group). C and F, were analyzed by Mann–Whitney U tests with unadjusted *P* values for the indicated comparisons; D, longitudinal blood measurements were analysed by repeated-measures two-way ANOVA with Tukey multiple-comparison testing, and terminal tissue measurements by repeated-measures two-way ANOVA with Bonferroni correction.; ns indicates not significant. Data are presented as mean ± SD. Statistical comparisons and sample sizes are indicated for each panel. Independent biological replicates represent individual mice.

To examine whether PE-MPs influence the relative contribution of *Tet2*-mutant haematopoiesis, we established a competitive bone marrow transplantation model. Irradiated male *Ldlr*^−/−^ mice received a 1:1 mixture of CD45.1 wild-type (WT) and CD45.2 *Tet2*^+/−^ donor marrow cells. Four weeks after transplantation, recipients were randomized to bilateral intratibial administration of 20-μm PE-MPs (1 μg per tibia) or phosphate-buffered saline, followed by 16 weeks of a high-fat, high-cholesterol diet (**Figure 1B**). The intratibial route and particle size were selected to investigate the consequences of local marrow exposure and retention rather than to reproduce systemic environmental exposure.

PE-MPs administration increased atherosclerotic lesion burden, as demonstrated by greater *en face* aortic arch plaque area and aortic-root Oil Red O-positive lesion area (**Figure 1C**). Concurrently, longitudinal flow cytometry showed a decline in the peripheral-blood CD45.1 WT/CD45.2 *Tet2*^+/−^ donor-cell ratio in both groups, with greater decline in PE-MPs-exposed mice from week 8 onward. At week 16, this ratio was also lower in the blood, bone marrow, spleen, and aorta of PE-MPs-exposed recipients (**Figure 1D**). These findings indicate that local marrow PE-MPs exposure increased atherosclerosis and shifted donor-cell representation towards *Tet2*^+/−^ haematopoiesis in this competitive transplantation model.

To explore the haematopoietic programmes associated with PE-MPs exposure, we performed bone marrow single-cell RNA sequencing in a separate cohort of irradiated *Ldlr*^−/−^ recipients reconstituted exclusively with *Tet2*^+/−^ marrow (PBS, n=3; PE-MPs, n=5). Reclustering of haematopoietic stem and multipotent progenitor cells (HSCs/MPPs) identified six transcriptional states, including cycling and activated HSCs and distinct MPP subsets (early MPP3, GMP-like MPP3, MPP2, and cycling MPP2). PE-MPs exposure was associated with granulocyte-monocyte lineage-priming and stress-response signatures, including expression of *Elane, Ctsg, Ly6c2, Jund*, and *Ier2* in selected HSCs/MPPs states. Regulon analysis further suggested increased activity of myeloid-associated transcriptional programmes involving *Spi1, Irf8*, and *Cebpb* (**Figure 1E**). These exploratory findings are consistent with altered myeloid-lineage programming following local PE-MPs exposure.

We then assessed *Tet2*^+/−^-derived myeloid populations in the competitive transplantation cohort. PE-MPs-exposed recipients exhibited higher absolute counts of *Tet2*^+/−^-derived granulocyte–monocyte progenitors in bone marrow, Ly6C^hi^ monocytes in peripheral blood, and Ly6C^hi^ macrophages populations in the spleen and aorta (**Figure 1F**). Together with the altered donor-cell ratios, these observations support increased relative representation and myeloid output of *Tet2*^+/−^ donor-derived haematopoiesis after PE-MPs exposure.

## Discussion

Our findings identify an association between local bone marrow PE-MPs exposure, altered competitive haematopoiesis, enhanced mutant-derived myeloid output, and accelerated atherosclerosis. The concurrent changes in donor-cell representation and lesion burden suggest a potential link between bone marrow microplastics and CH-associated vascular disease. However, the relative donor-cell ratios do not independently establish absolute clonal expansion, and the present model cannot determine whether the increased atherosclerosis is mediated by *Tet2*-mutant haematopoiesis or by parallel PE-MPs-induced effects. Moreover, direct intratibial administration of 20-μm particles does not recapitulate chronic systemic or local environmental exposure. These proof-of-concept findings warrant further investigation of whether environmentally relevant microplastic exposure modifies human CH and its cardiovascular consequences.

## Acknowledgements

The authors report the following contributions: conceptualization, Q.R.L., G.J.Q., X.W.; experiemnt design, X.N.G., Y.Y.L.; animal experiment, X.N.G., Y.Y.L.; clinical samples acquisition, X.W., X.N.G., Y.D.W.; data analysis, X.N.G., Y.Y.L., S.M.W.; consulting, H.B.Z.; writing (original draft), X.N.G.; writing (review and editing), Q.R.L., G.J.Q., X.W.; funding acquisition, Q.R.L., Y.Y.L., X.W.; supervision, Q.R.L., G.J.Q., X.W. This study was supported by grants from the National Natural Science Foundation of China (82270908, 82570558 to Q.R.L.; 82470464 to Y.Y.L.), Chongqing Science and Health Joint Medical Research Project (2023GDRC010) to X.W.

## References

1. Guo X, Wang L, Wang X, Li D, Wang H, Xu H, Liu Y, Kang R, Chen Q, Zheng L, Wu S, Guo Z, Zhang S. Discovery and analysis of microplastics in human bone marrow. Journal of Hazardous Materials 2024;477:135266.

2. Marfella R, Prattichizzo F, Sardu C, Fulgenzi G, Graciotti L, Spadoni T, D’Onofrio N, Scisciola L, La Grotta R, Frigé C, Pellegrini V, Municinò M, Siniscalchi M, Spinetti F, Vigliotti G, Vecchione C, Carrizzo A, Accarino G, Squillante A, Spaziano G, Mirra D, Esposito R, Altieri S, Falco G, Fenti A, Galoppo S, Canzano S, Sasso FC, Matacchione G, Olivieri F, Ferraraccio F, Panarese I, Paolisso P, Barbato E, Lubritto C, Balestrieri ML, Mauro C, Caballero AE, Rajagopalan S, Ceriello A, D’Agostino B, Iovino P, Paolisso G. Microplastics and Nanoplastics in Atheromas and Cardiovascular Events. The New England Journal of Medicine 2024;390(10):900–910.

3. Schuermans A, Honigberg MC. Clonal haematopoiesis in cardiovascular disease: prognostic role and novel therapeutic target. Nature Reviews. Cardiology 2025;22(11):845–856.

4. Libby P, Sidlow R, Lin AE, Gupta D, Jones LW, Moslehi J, Zeiher A, Jaiswal S, Schulz C, Blankstein R, Bolton KL, Steensma D, Levine RL, Ebert BL. Clonal Hematopoiesis: Crossroads of Aging, Cardiovascular Disease, and Cancer: JACC Review Topic of the Week. Journal of the American College of Cardiology 2019;74(4):567–577.

5. Fuster JJ, MacLauchlan S, Zuriaga MA, Polackal MN, Ostriker AC, Chakraborty R, Wu CL, Sano S, Muralidharan S, Rius C, Vuong J, Jacob S, Muralidhar V, Robertson AA, Cooper MA, Andrés V, Hirschi KK, Martin KA, Walsh K. Clonal hematopoiesis associated with TET2 deficiency accelerates atherosclerosis development in mice. Science 2017;355(6327):842–847.

